# Non-apoptotic enteroblast-specific role of the initiator caspase Dronc for development and homeostasis of the *Drosophila* intestine

**DOI:** 10.1101/2020.10.25.354431

**Authors:** Jillian L. Lindblad, Meghana Tare, Alla Amcheslavsky, Alicia Shields, Andreas Bergmann

## Abstract

The initiator caspase Dronc is the only CARD-domain containing caspase in *Drosophila* and is essential for apoptosis. Here, we report that homozygous *dronc* mutant adult animals are short-lived due to the presence of a poorly developed, defective and leaky intestine. Interestingly, this mutant phenotype can be significantly rescued by enteroblast-specific expression of *dronc*^+^ in *dronc* mutant animals, suggesting that proper Dronc function specifically in enteroblasts, one of four cell types in the intestine, is critical for normal development of the intestine. Furthermore, enteroblast-specific knockdown of *dronc* in adult intestines triggers hyperplasia and differentiation defects. These enteroblast-specific functions of Dronc do not require the apoptotic pathway and thus occur in a non-apoptotic manner. In summary, we demonstrate that an apoptotic initiator caspase has a very critical non-apoptotic function for normal development and for the control of the cell lineage in the adult midgut and therefore for proper physiology and homeostasis.

**Highlights:**

- *dronc* mutants die from a fragile and leaky intestine
- *dronc* has a critical function in enteroblasts of the intestine
- *dronc* controls proliferation and differentiation in the intestine
- *dronc* performs these functions in an apoptosis-independent (non-apoptotic) manner

## Introduction

Caspases are important executioners of programmed cell death (apoptosis) in multicellular organisms. They encode highly specific Cys proteases which are produced as inactive zymogens in living cells. There are two different types of caspases. Initiator caspases (Caspase-8, Caspase-9 and *Drosophila* Dronc) act upstream of effector caspases (Caspase-3, Caspase-7 and *Drosophila* DrICE) ^1,2^. Initiator caspases are distinguished from effector caspases by the presence of long prodomains that harbour protein/protein interaction domains such as the CARD (caspase activation and recruitment domain) in Caspase-9 and Dronc. In *Drosophila*, activation of Dronc proceeds through CARD/CARD interactions with the adaptor protein Dark ^3–6^ which leads to formation of the apoptosome ^7–9^ In the apoptosome, Dronc cleaves and activates effector caspases such as DrICE ^1,2,10^. After activation, these effector caspases cleave a large number of cellular proteins initiating the demise of the cell.

Homozygous *dronc* mutants are strongly semi-lethal. Most of them die during pupal development, but at a very low frequency (< 1% of the expected offspring) homozygous escaper flies can be recovered. These escapers exhibit down-curved, opaque wings ^11^. This phenotype is likely the result of failed apoptosis during wing maturation ^11^. Other than the wing phenotype, any other obvious phenotype of homozygous adult *dronc* mutants have not been reported. However, they do have a very short life-span. They only live for about 3-4 days after eclosion. The cause for this short life-span is not known and will be examined in this study.

In addition to the apoptotic function of caspases, there is a growing list of non-apoptotic functions in basic cellular processes such as proliferation, differentiation, cell migration, sperm maturation and others ^12–14^. When these processes require non-apoptotic caspase function, they are also referred to as caspase-dependent non-lethal cellular processes (CDPs) ^15^. Very sensitive reporters of caspase activity have revealed that many cells experience caspase activity at one point in their life without triggering apoptosis ^16–18^. How cells escape the potentially cell lethal activity of active caspases is currently subject of intense research.

The *Drosophila* midgut has emerged as a valuable model to study adult stem cells. It is composed of only 4 different cell types: intestinal stem cells (ISC), enteroblasts (EB), enterocytes (EC) and secretory enteroendocrine cells (EE) ^19,20^ (reviewed in ^21^). The ISCs are the only proliferating cells in the midgut. EBs are produced by asymmetric division of ISCs and are transient cells which differentiate into ECs or EEs ^19,20^. ECs are absorptive epithelial cells and make up the majority of the cells in the midgut. More recently, it has also been suggested that EEs can directly emerge from ISC division without going through an EB intermediate ^22,23^.

The cell lineage in the midgut is under strict control to ensure proper function and homeostasis of the midgut (reviewed in ^21,24^ ECs and EEs are regularly turned over and need to be replaced by new cells due to ISC mitosis. There is feedback from dying EC cells to control ISC activity ^25^. Imbalances of this control can lead to malfunction of the intestine, dysplasia, premature aging and death of the animal. Because mammalian intestines are also subject to a similar cell lineage ^26^, a clear understanding of the mechanisms involved in the control of this cell lineage may help to understand disease and identify potential targets for the cure of the disease.

Here, we show that the initiator caspase Dronc has essential functions for development and homeostasis of the *Drosophila* adult midgut. Interestingly, this function is primarily required in EBs and appears to be non-apoptotic in nature. Loss of *dronc* in EBs causes hyperplasia due to increased proliferation. Furthermore, there are significant differentiation defects. Specifically, loss of *dronc* results in an increased number of EBs and accumulation of cells that have features of both EBs and ECs. There is also a significant increase in the number of EEs. These data demonstrate that an apoptotic initiator caspase has a very critical function for the control of the cell lineage in the adult midgut and therefore for proper physiology and homeostasis.

## Results

### Homozygous *dronc* mutants die prematurely due to fragile and leaky intestines

Homozygous *dronc* mutants are pupal lethal. However, at a very low frequency (less than 1% of the expected offspring), adult homozygous *dronc*^*I24/I29*^ mutant flies can be recovered. These *dronc* alleles carry premature stop codons at codons 28 and 53 and encode strong, if not null, alleles of *dronc* ^11^. With the exception of down-curved and opaque wings ^11^, these mutant flies do not have any obvious phenotypic abnormalities. Nevertheless, they are very short-lived and die within 3 to 4 days after eclosion suggesting that they may have some internal defects. To identify the possible cause of this premature death, we examined the internal organs of these mutants. When dissecting the intestines of *dronc*^*I24/I29*^ flies, we noticed that they are very fragile. By phallodin labelings, these intestines displayed structural irregularities (Figure 1A,B). Furthermore, labelings with the nuclear dye DAPI show that the mutant intestines have a higher density of cells (Figure 1A’,B’; quantified in 1C).

**Figure 1.**
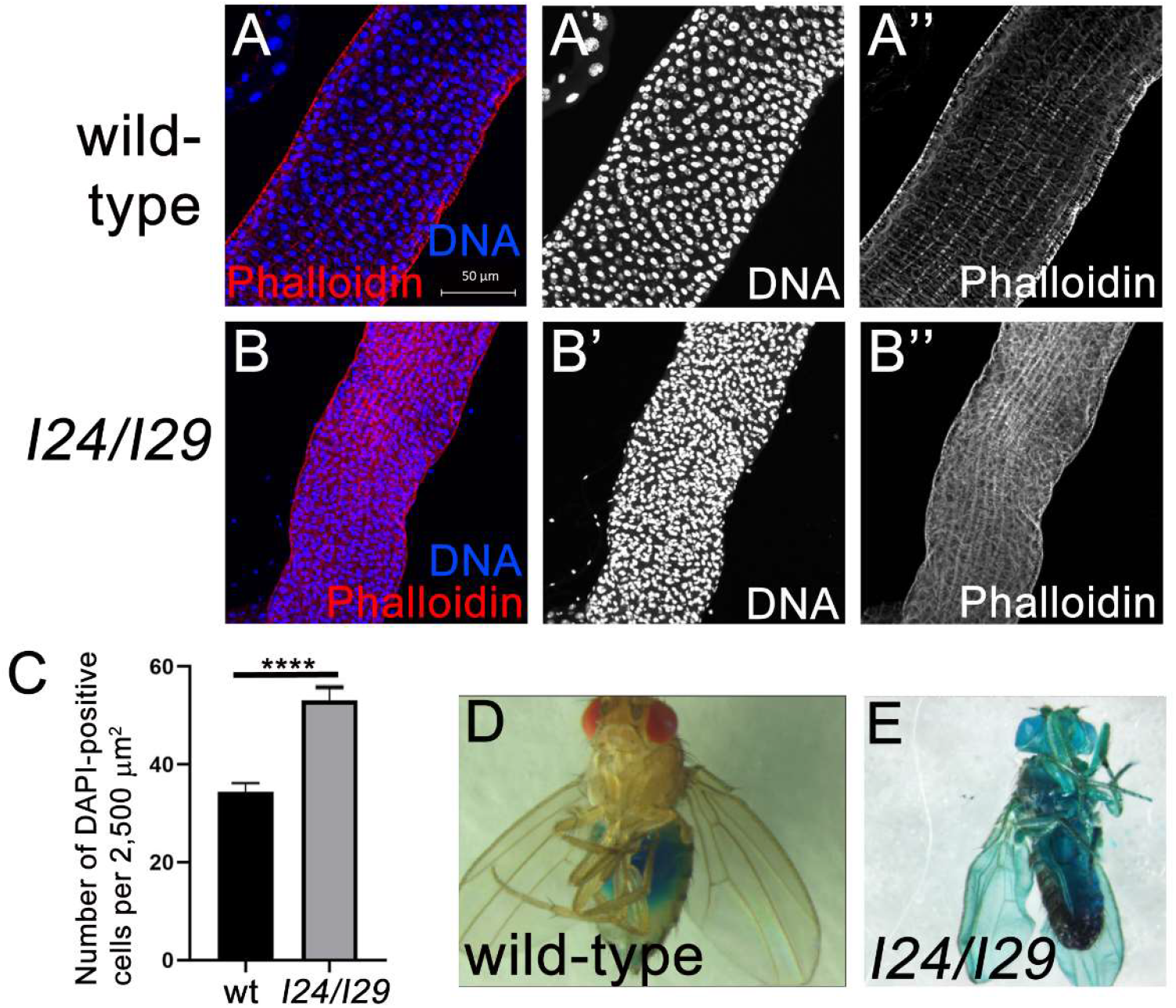
Homozygous *dronc*^*I24/*^*dronc*^*I29*^ mutants have defective and leaky intestines. **(A-C)** DAPI- and Phalloidin-labeled wild-type (Canton S) and *dronc*^*I24*^/*dronc*^*I29*^ *(124/129)* intestines. The examples shown are taken from Region R4ab, but are representative for the entire intestine. Scale bar: 50 μm. (C) Quantification of the DAPI labelings in wt and *dronc*^*I24*^/*dronc*^*I29*^ intestines. The density of nuclei in 2,500 μm^2^ fields was analysed by unpaired t test, two tailored and plotted± SEM. ****p < 0.0001. n = 6 (wt), 5 (*dronc*^*I24*^/*dronc*^*I29*^). **(F,G)** Wild-type (Canton S) and *dronc*^*I24/I29*^ mutant animals, subjected to a SMURF assay.

To examine if these structural defects cause a dysfunction of the intestine, we performed SMURF assays with homozygous *dronc*^*I24/I29*^ flies. In a SMURF assay, a blue dye is mixed into the food and fed to the flies ^27,28^. Flies with an intact intestine keep the blue food in the intestine which can be easily seen through the abdominal cuticle (Figure 1D). However, in flies with an intestinal barrier dysfunction, the blue dye penetrates into every tissue of the fly, generating a SMURF phenotype ^28^. We examined five homozygous *dronc*^*I24/I29*^ mutants in the SMURF assay and all of them displayed the SMURF phenotype (Figure 1E). Furthermore, while wild-type flies start feeding almost immediately, we noticed that *dronc*^*I24/I29*^ mutant flies do not feed for the first 24 to 48h after eclosion. These data suggest that *dronc* mutant intestines have structural defects and a defective barrier function causing a leaky gut.

### EB-specific expression of *dronc* rescues semi-lethality and restores gut function of *dronc* mutants

To determine if the defective intestine causes the premature organismal lethality of homozygous *dronc*^*I24/I29*^ mutants, we asked if cell-type specific expression of *UAS-dronc*^*wt*^ can rescue the strong semi-lethality and short lifespan as well as the defective and leaky gut phenotype of homozygous *dronc*^*I24/I29*^ animals. As a control, we expressed the catalytic *UAS-dronc*^*C318A*^ mutant. Interestingly, expression of *dronc*^*wt*^ in EBs using the EB-specific driver *Su(H)GBE-Ga/4* (from now on *Su(H))* gave the best rescue of the lethality (Figure 2A; genotype 3). About 75% of the expected *Su(H)>dronc*^wt^; *dronc*^*I24/I29*^ progeny was recovered as adults. With the *esg-Gal4* driver, which is expressed in ISCs and to some extent in EBs, a weaker rescue was recorded (Figure 2A; genotype 2). The weakest rescue was scored when *dronc*^*wt*^ was expressed in mature ECs using *NP1-Gal4* (Figure 2A; genotype 4). Expression of the catalytic mutant *dronc*^*C318A*^ using all three Gal4 drivers was not able to rescue the lethality of *dronc*^*I24/I29*^ animals (Figure 2A; genotypes 5-7). These rescue crosses suggest that for development of the intestine, Dronc function is most critical in EBs and requires its catalytic activity.

**Figure 2.**
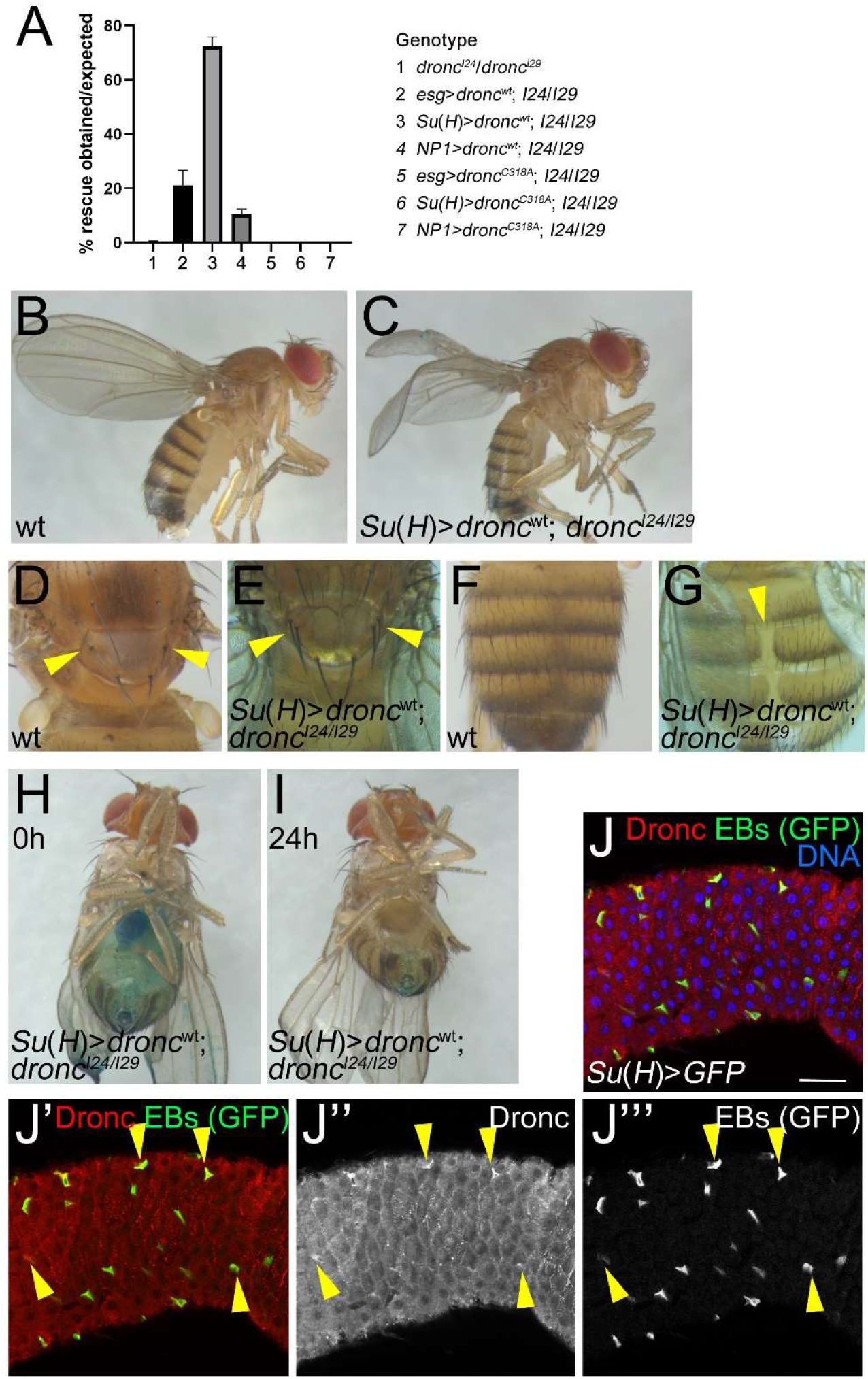
Enteroblast-specific expression of *dronc*^*wt*^ rescues the semi-lethality and leakage of homozygous *dronc* mutant animals. **(A)** *esg-Gal4* (expression in ISCs), *Su(H)-Gal4* (EBs) and *NP1-Gal4* (ECs) were used to drive expression of *UAS-dronc*^*wt*^ or *UAS-dronc*^*C318A*^ in *dronc*^*I24*^/*dronc*^*I29*^ *(I24/I29)* mutants and survival was scored. *dronc*^*C318A*^ carries a mutation of the catalytic Cys318 residue to an Ala and represents a catalytic mutant. *dronc*^*I24/I29*^ (genotype 1) is strongly pupal semi-lethal and less than 1% of the expected progeny can be recovered as adults. The obtained progeny in the rescue crosses is plotted as the percentage of the expected Mendelian progeny. Genotypes of expected rescued progeny are indicated on the right. The average of three independent experiments is shown. At least 200 flies were counted in each experiment. Rescue data were analyzed by one-way ANOVA with Holm-Sidak test for multiple comparisons. Error bars are SEM. **(B,C)** While the wmgs of wild-type (Canton S) flies are straight (B), the wmgs of *Su(H)>dronc*^wt^; *dronc*^*I24/I29*^ rescued flies are down-curved (C) similar to *dronc*^*I24/I29*^ mutants^11^ **(D-G)** *Su(H)>dronc*^wt^; *dronc*^*I24/I29*^ flies exhibit morphological phenotypes. Canton S (wt) flies contain 4 scutellar bristles (macrochaetae) (D) and have a closed abdomen (F). All *Su(H)>dronc*^wt^; *dronc*^*I24/I29*^ rescued flies have duplications of 1 or 2 macrochaetae (D, arrowheads) and about 50% display a split abdomen phenotype (G, arrowhead) (n>50). **(H,I)** 25 out of 26 tested *Su(H)>dronc*^wt^; *dronc*^*I24/I29*^ show a wild-type SMURF phenotype (H) (compare to Figure 1D,E). 24h after removal from blue food, these flies have completely cleared the gut of blue food (I). **(J)** Labeling of *Su(H)>GFP* midguts with anti-Dronc antibodies (red in J and J’; grey in J’’). EBs are labelled by GFP (green in J and J’; grey in J’’’). Yellow arrows highlight several EBs containing Dronc protein. Scale bar: 50 μm.

However, these data have the caveat that the Gal4 drivers used are also expressed in other tissues during development. We can therefore not conclude, that the exclusive expression of *dronc* in EBs is sufficient for normal development and survival. Nevertheless, EB-specific expression of *dronc* does not rescue all known phenotypes of *dronc* mutants. The wings of *Su(H)>dronc*^wt^; *dronc*^*I24/I29*^ flies still have the reported *dronc* mutant phenotype (down-curved, opaque wings) (Figure 2B,C). Furthermore, we observed one to two additional scutellar bristles (macrochaetae) with 100% penetrance (Figure 2D,E) which had been reported for *dark* and *cytochrome c-d* mutants ^29,30^ and represents a typical phenotype when cell death is blocked. We also recovered *Su(H)>dronc*^wt^; *dronc*^*I24/I29*^ flies with a split abdomen phenotype at reduced penetrance (Figure 2F,G). The split abdomen phenotype is likely the result of reduced cell death of larval cells in the abdomen during pupal development ^31^. Therefore, despite the caveat that *Su(H)-Gal4* expression is not restricted to EBs in the midgut, Su(H)-driven expression of *dronc* cannot rescue all phenotypes of *dronc*^*I24/I29*^ mutants, demonstrating that expression of *dronc* in select groups of cells is sufficient for development and survival of the animal.

To directly examine the effect of EB-specific expression of *dronc* on the physiology of the intestine, we performed SMURF assays with *Su(H)>dronc*^wt^; *dronc*^*I24/I29*^ rescued animals. Of 26 rescued flies which were tested in this assay, only one animal developed a SMURF phenotype. The intestines of the other 25 flies stayed intact and were also able to clear the blue food within 24 hours after they were removed from the blue food (Figure 2H,I). In contrast, 6 out of 7 tested *esg>dronc*^wt^;*dronc*^*I24/I29*^ flies (Figure 2A, genotype 2) developed a SMURF phenotype.

Although we did not perform extended life span assays, *Su(H)>dronc*^wt^; *dronc*^*I24/I29*^ flies lived significantly longer(> 2 weeks) than *dronc*^*I24/I29*^ flies which live only for 3-4 days after eclosion. The life span of *esg>dronc*^wt^; *dronc*^*I24/I29*^ also improved, but most of them died after about 1 week.

Given the critical role of Dronc in the adult intestine, in particular in EBs, we examined if Dronc is expressed in these cells using Dronc-specific antibodies. To identify EBs, we expressed GFP using the EB-specific driver *Su(H)-Gal4*. This analysis reveals that Dronc is - among other cells - expressed in EBs (Figure 2J, yellow arrowheads). Taken together, these results indicate that Dronc function in the intestine, in particular in EBs, is essential for intestinal integrity and organismal survival of the animal.

### Down-regulation of *dronc* in EB cells causes hyperplasia

The above data indicate that *dronc* has a very important function in EB cells for proper formation of the intestine during development. Additionally, we examined if *dronc* also has an important role for homeostasis of the adult midgut. For that purpose, we down-regulated *dronc* in EB cells by RNAi, using *Su(H)-Gal4*^*ts*^. The ^*ts*^ in this annotation indicates the presence of *Gal80*^*ts*^ which allows down-regulation of *dronc* using this Gal4 driver by temperature shift to 29°C after the animals have fully developed and eclosed. 5 days old *Su(H)*^*ts*^>*dronc*^*RNAi*^ flies were shifted from 18°C to 29°C and their intestines were analysed 5-6 days later. Preferentially, region R4ab of the posterior midgut was examined in these assays ^32,33^ There is a significant increase in the total number of cells in the midgut (as revealed by DAPI labelings (Figure 3A-C’’, quantified in 3D)), and GFP-positive EB cells were overabundant compared to controls (Figure 3A’-C’, quantified in 3E). Consistent results were obtained for 2 independent *dronc* RNAi lines (Figure 3B-E). In addition to the EB overabundance of *Su(H)*^*ts*^>*dronc*^*RNAi*^ midguts, EBs (marked by GFP) also appear to change shape and are much larger than normal (Figure 3A’-C’). This observation is also statistically significant (Figure 3F).

**Figure 3.**
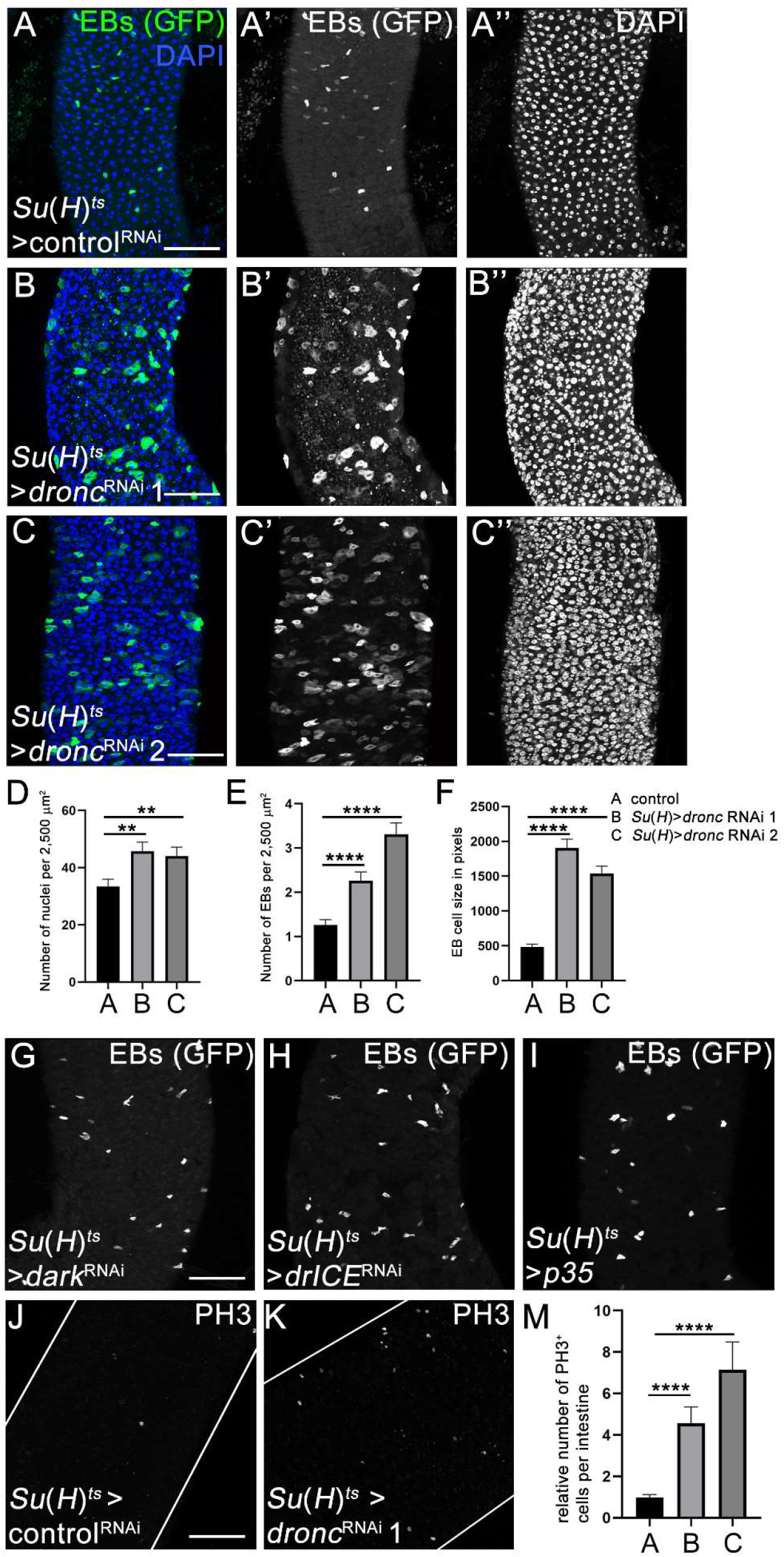
EB-specific knockdown of *dronc* causes accumulation of EBs and hyperplasia. In this and the following figure, crosses were incubated at 18°C until adults eclosed. 5 days old females of the indicated genotype were shifted to 29°C and incubated for 5-6 days until dissection and labelling of the midguts. Two different *dronc* RNAi transgenes were used: line 1 = P{GD12376}v23033; line 2 = P{KK104278}v100424. *Lueiferase* RNAi was used as a control in (A). **(A-C)** Control (A) and *Su(H)*^*ts*^>*dronc* RNAi 1 and 2 (B,C) midguts were labelled with GFP (green in A-C; grey in A’ -C’) and DAPI (blue in A-C; grey in A’’ -C’’). Note that in addition to an increased number of EBs in response to *dronc* RNAi, these cells also underwent morphological changes and appear much larger compared to control (A). Scale bar: 50 μm. Complete genotypes: (A) *Su(H)-Gal4 UAS-GFP tub-Gal80*^*ts*^; *UAS-lueiferase* RNAi; (B) *Su(H)-Gal4 UAS-GFP tub-Gal80*^*ts*^; *UAS-dronc* RNAi 1; (C) *Su(H)-Gal4 UAS-GFP tub-Gal80*^*ts*^; *UAS-dronc* RNAi 2. **(D,E)** Quantification of the numbers of nuclei (DAPI) and EBs (GFP) of the genotypes in (A­ C). To obtain DAPI counts, at least five random, but representative fields of 2,500 μm^2^ each per posterior midgut were counted. In the case of EBs, all GFP-positive cells (corresponding to EBs) were counted in region R4ab and normalized to 2,500 μm^2^ fields. Data were analyzed by one-way ANOVA with Holm-Sidak test for multiple comparisons. Error bars are SEM. ****p < 0.0001; **p< 0.01. Number of midguts analysed: n=12 (control), 9 (*dronc* RNAi 1), 12 (*dronc* RNAi 2). **(F)** Quantification of the cell size of EBs of control and *dronc* RNAi midguts. Shape of EBs was outlined using the quick selection tool in Photoshop and the number of pixels in the selected area was recorded. Data were analyzed by one-way ANOVA with Holm-Sidak test for multiple comparisons. Error bars are SEM. ****p < 0.0001. n=24 (control), 25 (*dronc* RNAi 1), 24 (*dronc* RNAi 2). **(G-1)** EB-specific knockdown of *dark* (G) and *drICE* (H) by RNAi or overexpression of *p35* (I) does not phenocopy the phenotypes observed by *dronc* RNAi (compare to (B,C)). More examples and quantifications are shown in Supplemental Figure S1. EBs were labelled with GFP. The conditions were the same as in (A-C). Scale bars: 50 μm. Complete genotypes: (G) *Su(H)-Gal4 UAS-GFP tub-Gal80*^*ts*^; *UAS-dark* RNAi; (H) *Su(H)-Gal4 UAS-GFP tub-Gal80*^*ts*^; *UAS-drICE* RNAi; (I) *Su(H)-Gal4 UAS-GFP tub-Gal80*^*ts*^; *UAS-p35*. **(J,K)** PH3 labelings of control and *Su(H)*^*ts*^>*dronc* RNAi midguts. Scale bar: 50 μm. Complete genotypes: (A) *Su(H)-Gal4 UAS-GFP tub-Gal80*^*ts*^; *UAS-lueiferase* RNAi (B) *Su(H)-Gal4 UAS-GFP tub-Gal80*^*ts*^; *UAS-dronc* RNAi (P{GD12376}v23033). **(M)** Quantification of PH3 labelings of control and *Su(H)*^*ts*^>*dronc* RNAi midguts. PH3 cells were manually counted across the entire intestine and analysed by one-way ANOVA with Holm-Sidak test for multiple comparisons. Error bars are SEM. ****p < 0.0001. n=l 0 (control), 11 (*dronc* RNAi 1), 10 (*dronc* RNAi 2). Genotypes as in (A-C).

Given that *dronc* has an important function in apoptosis ^11,34–36^, we considered the possibility that this EB-specific *dronc* phenotype is caused by loss of apoptosis. However, we did not observe a similar EB-overabundance phenotype in response to EB-specific (using *Su(H)-Gal4*^*ts*^) *dark* RNAi (Figure 3G; see additional example and quantification in Supplementary Figure S1), which encodes the adaptor protein for incorporation of Dronc into the apoptosome during apoptosis ^3,4,6^. EB-specific RNAi targeting *drICE*, the most important effector caspase in *Drosophila* ^37,38^, also did not phenocopy the *dronc* RNAi phenotype (Figure 3H; Supplementary Figure S1). Finally, EB-specific expression of the effector caspase inhibitor *p35* in otherwise wild-type midguts using *Su(H)-Gal4*^*ts*^ did not replicate the *dronc* RNAi phenotype (Figure 3I; Supplementary Figure S1). EB-specific *dark*^RNAi^, *drICE*^RNAi^ and *p35* expression also did not phenocopy other aspects of the *dronc*^RNAi^ phenotype such as the increase in overall cell number and in EB cell size (Supplementary Figure S1). Therefore, we can rule out that the *Su(H)*^*ts*^>*dronc*^*RNAi*^ phenotypes in adult midguts are caused entirely by loss of apoptosis. The functionality of the *dark*^RNAi^, *drICE*^RNAi^ and *p35* transgenic lines was validated by the ability of these lines to suppress the *GMR-reaper* eye ablation phenotype, a commonly used apoptosis model ^39^ (Supplementary Figure S1).

It was previously shown that an accumulation of EBs can cause increased ISC proliferation ^40^. Consistently, based on PH3-labeling experiments, we found that cell proliferation is increased in *Su(H)*^*ts*^>*dronc*^*RNAi*^ midguts (Figure 3J,K). This was consistent with two *dronc*^RNAi^ lines (Figure 3M). In contrast, EB-specific *dark*^RNAi^, *drICE*^RNAi^ and *p35* expression did not phenocopy this phenotype (Supplementary Figure S1) suggesting that *dronc* controls cell proliferation in a non-apoptotic manner. Combined, these observations suggest that EB-specific *dronc* RNAi triggers a hyperplastic phenotype in the intestine.

### Down-regulation of *dronc* in EB cells causes differentiation defects

Given that ECs are much larger in size than EBs and that EBs in *Su(H)>dronc*^RNAi^ midguts are significantly enlarged (Figure 3F), we considered that GFP-positive EBs in *dronc*^RNAi^ midguts also display features of EC cell fate. To examine this possibility, we labelled these midguts with Pdm-1 antibody, a marker for EC fate ^41,42^ Consistently, in *Su(H)*^*ts*^>*dronc*^*RNAi*^, *GFP* midguts we observed multiple examples where *Su(H)*^*ts*^-driven GFP overlaps with Pdm-1 labeling suggesting that these cells have properties of both EBs and ECs (Figure 4A-C). While in control midguts, expression of Pdm-1 in EBs was also observed at low frequency (~2% of total EBs), this number was significantly higher in *Su(H)*^*ts*^>*dronc*^*RNAi*^ midguts (Figure 4G). These data imply that in normal midguts, Dronc is required for the appropriate differentiation of EBs into ECs.

**Figure 4.**
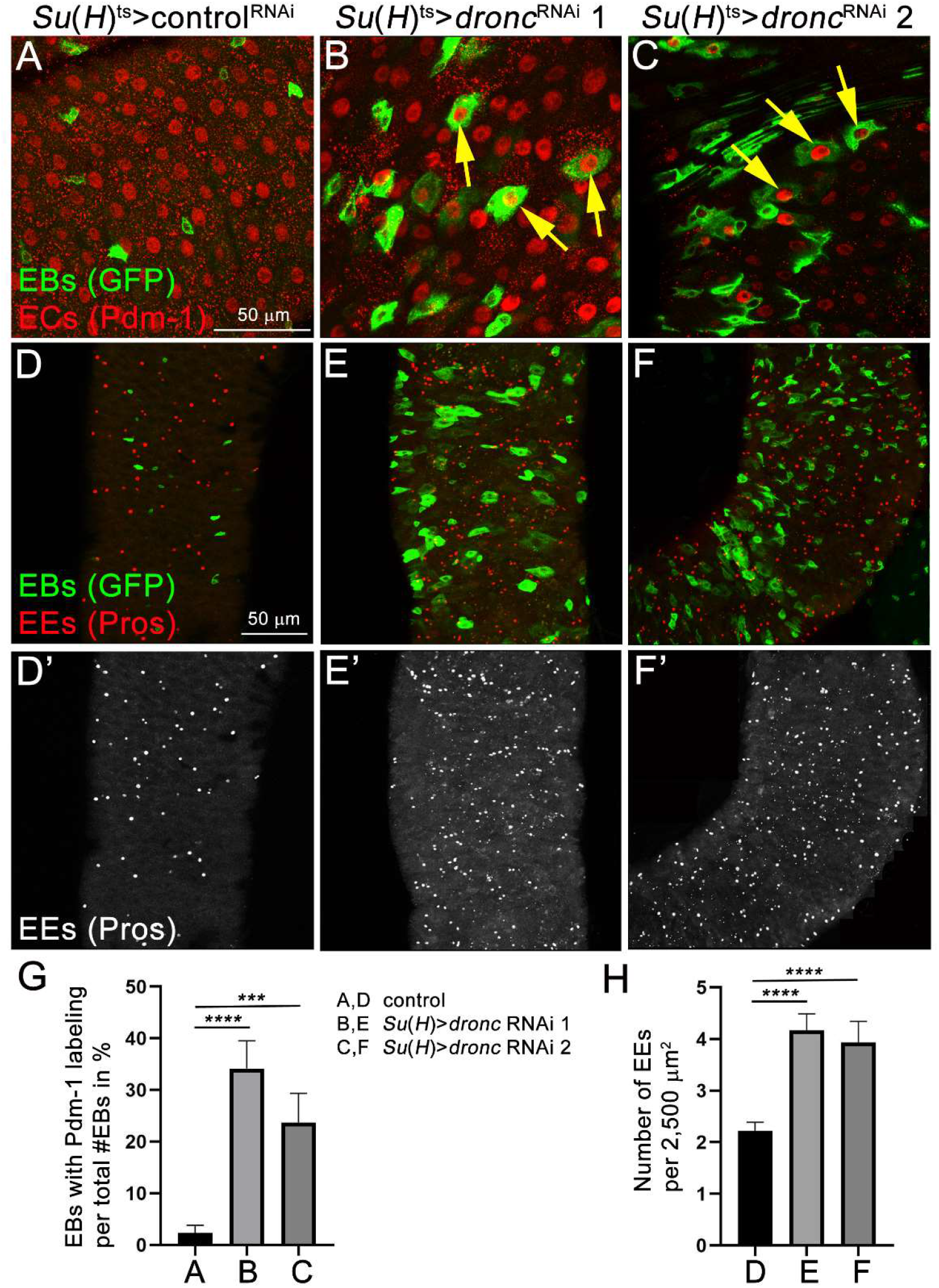
Down-regulation of *dronc* in EBs causes differentiation defects. **(A-C)** Shown are regions R4ab of control midguts (A) and of midguts in which *dronc* was down-regulated in EB cells using *Su(H)-Gal4*^*ts*^ (B,C). These midguts were labelled with GFP (to visualize EBs) and Pdm-1 antibodies which labels nuclei of ECs. Yellow arrows in (B,C) point to GFP/Pdm-1 double positive cells. Two different *dronc* RNAi transgenes were used: line 1 = P{GD12376}v23033; line 2 = P{KK104278}v100424. *Luciferase* RNAi was used as a control in (A). Complete genotypes: (A) *Su(H)-Gal4 UAS-GFP tub-Gal80*^*ts*^; *UAS-luciferase* RNAi; (B,C) *Su(H)-Gal4 UAS-GFP tub-Gal80*^*ts*^/*UAS-dronc* RNAi line 1 (B)/line 2 (C). Scale bar: 50 μm. **(D-F)** Prospero (Pros) antibody (red in D-F; grey in D’-F’) was used to label enteroendocrine cells (EE) in regions R4ab of control midguts (D) and midguts in which *dronc* was down-regulated in EBs using *Su(H)-Gal4*^*ts*^ (E,F). EBs are labelled by GFP (green in D-F) due to *Su(H)>GFP* expression. Genotypes as in (A-C). Scale bar: 50 μm. **(G)** Quantification of the data in (A-C). GFP/Pdm-1 double positive cells were normalized to the total number of GFP-positive cells and illustrated in %. GFP/Pdm-1 double positive and GFP-positive cells were manually counted and analysed by one-way ANOVA with Holm-Sidak test for multiple comparisons. Error bars are SEM. ****p < 0.0001; ***p < 0.001. Number of midguts analysed: n=7 (control), 4 (*dronc* RNAi 1), 4 (*dronc* RNAi 2). **(h)** Quantification of the Pros labelings in (D’-F’). All Pros positive cells (corresponding to EEs) were counted in region R4ab and normalized to 2,500 μm^2^ fields. Data were analyzed by one-way ANOVA with Holm-Sidak test for multiple comparisons. Error bars are SEM. ****p < 0.0001. Number of midguts analysed: n=12 (control), 7 (*dronc* RNAi 1), 6 (*dronc* RNAi 2).

We also examined enteroendocrine (EE) cells in *Su(H)*^*ts*^>*dronc*^*RNAi*^ midguts using the EE marker Prospero (Pros) and observed an increased number of Pros-positive cells (Figure 4D’-F’; quantified in 4H), indicating that the number of EEs is elevated. This EE overabundance phenotype was not observed in response to EB-specific *dark*^RNAi^, *drICE*^RNAi^ and *p35* expression in adult midguts (Supplementary Figure S1) suggesting that this *dronc*^RNAi^ dependent phenotype is independent on its role in apoptosis. These observations further indicate that Dronc function in EBs is required for proper differentiation of intestinal cell types.

## Discussion

In the absence of *dronc* during development, the adult intestine has structural defects, is fragile and leaky, limiting the lifespan of homozygous adult escapers to only 3-4 days. Interestingly, expression of *dronc* specifically in EBs can significantly rescue these phenotypes suggesting that *dronc* has a very important function in EBs for development of the intestine. Because the catalytic mutant *dronc*^*C318A*^ cannot rescue these phenotypes, Dronc likely requires its catalytic activity for proper function in the midgut. The observation that expression of *dronc* in ECs using *NPI-Gal4* cannot significantly rescue the lethality of *dronc* mutants (Figure 2A) does not mean that Dronc does not have a function in ECs. It only means that *NPI-Gal4* is not expressed in those cells (such as EBs) where *dronc* has an essential function for survival.

EB-specific knockdown of *dronc* in adult intestines causes differentiation defects and hyperplasia due to increased proliferation. Therefore, Dronc function is critical for proper control of the cell lineage in the midgut and loss of *dronc* disrupts this homeostasis. The observation that loss of other genes in the apoptosis pathway *(dark, drICE)* or overexpression of the effector caspase inhibitor *p35* do not replicate the EB-specific *dronc* phenotypes suggests that Dronc mediates this role in a non-apoptotic manner. This adds control of proper proliferation and differentiation of the adult midgut to the growing list of non-apoptotic functions of Dronc.

There are several interesting aspects of the EB-specific *dronc* phenotypes in the midgut. First, the observed hyperplasia is puzzling. In the *Drosophila* intestine, only ISCs are mitotically active, EBs are not ^19,20^. In fact, EBs are the daughter cells of the asymmetric division of ISCs. The increased mitotic activity of ISCs in response to EB-specific *dronc* knockdown suggests that Dronc is involved in a feedback mechanism between EBs and ISCs. Elucidating the molecular mechanism of this feedback mechanism and the role Dronc plays in this will be an exciting avenue for future research.

Second, the increased number of EBs and accumulation of EBs expressing the EC marker Pdm-1 suggests that Dronc is involved in the differentiation process from EBs to ECs. The function of Dronc in this process can be explained in one of two opposite ways. Dronc may be required for an important step in the differentiation process from EBs to ECs. In the absence of *dronc*, while EBs are able to increase in size and induce expression of the EC marker Pdm-1, they can no longer complete the differentiation program into ECs and get stuck along the way. This would result in an increased number of EBs in the intestine. However, the opposite explanation, that Dronc inhibits the differentiation of EBs into ECs under normal conditions, is also formally possible. In that case, the differentiation into ECs occurs so fast, that the EB-specific expression of GFP (which is *Su(H)-Gal4* dependent) is not turned off early enough to avoid overlap of EB- and EC-specific markers. Future experiments will clarify by which mechanism Dronc controls the differentiation of EBs to ECs.

Third, the strong increase of EEs in response to EB-specific knockdown of *dronc* suggests that Dronc negatively controls the formation and differentiation of EEs. Because it is not clear whether EEs are also differentiating from EBs, as originally suggested ^19,43^, or if they are direct descendants of ISCs ^22,23^, it is not clear whether the role of Dronc in this process is autonomous or non-cell autonomous. Because we do not observe an overlap of EB-specific GFP expression (driven by *Su(H)-Gal4)* with EE-specific Pros labelling (Figure 4D-F), suggest a non-cell autonomous control of EE fate by Dronc, but other explanations may be possible, too. In any case, what these data show is that under normal conditions, Dronc controls the number of EEs in an EB-specific manner.

Another important question for the future is how Dronc mediates the homeostatic effect in the adult intestine. There are several possibilities. It was shown that loss of the transcription factor *eseargot (ese)* has similar phenotypes compared to *dronc:* increased differentiation into ECs and EEs ^44,45^. Esc suppresses the expression of the differentiation-promoting factor Pdm-1 in progenitor cells ^44^. Thus, Dronc may be involved in Esc-mediated control of Pdm-1 expression. Dronc may also be involved in the control of some of the signalling pathways that operate in the differentiation process, such as Notch (N) signalling. Given that N signalling is also controlled by *esc* ^44,45^, it is possible that Dronc participates in this complex signalling network. However, because many signalling pathways are involved in the control of the cell lineage in the intestine (reviewed in ^21,24^), Dronc may also control any of these pathways. Given that the only known biochemical function of Dronc is proteolytic activity, it will be a challenge in the future to identify the cleavage substrate(s) of Dronc in this process.

Finally, how is Dronc activated in this non-apoptotic context? During apoptosis, activation of Dronc occurs by incorporation into the Dark apoptosome ^7–9^ However, in EBs of the intestine, this occurs in an apoptosome-independent manner because Dark is not involved (Figure 3G; Supplementary Figure S1). Dronc may be incorporated into a different protein complex for activation. For example, mammalian initiator caspases can be recruited into different complexes such as the apoptosome and the inflammosome ^46^. A different complex may provide different properties to Dronc such that it does not cleave its apoptotic substrates and can act in a non-apoptotic manner. Alternatively, it is possible that a different protease may cleave and activate Dronc.

Impaired caspase function may also provide a contributing factor for the development of colon cancer in human patients. For example, Caspase-9, the Dronc ortholog in humans, is epigenetically silenced in almost 50% of colon cancer cases ^47,48^. Other caspases, such as Caspase-7 are down-regulated in up to 85% of colon cancer cases ^47,48^. While this silencing is likely a means for evading apoptosis, it could also trigger additional effects such as increased proliferation and differentiation defects, thus further supporting tumorigenesis. Therefore, revealing the mechanism by which Dronc expression in EBs maintains tissue homeostasis may also have important implications for understanding of the tumor-promoting effect of loss of caspase-9 in humans. Given that the function of Dronc for maintaining tissue homeostasis in the intestine is non-apoptotic, characterizing the role of Dronc in EBs does provide a convenient opportunity to study this function in the absence of its apoptotic function which may not be as simple for Caspase-9 in humans.

## Resource Availability

### Materials Availability

All materials and fly stocks used in this study are available from the corresponding author on request.

### Data Availability

All data generated or analysed during this study are included in this published article (and its Supplementary Information files).

## Acknowledgements

We would like to thank Yu Cai, Fernando Diaz-Benjumea, Tony Ip, Pascal Meier, the Bloomington *Drosophila* Stock Center (BDSC), the Vienna *Drosophila* Resource Center (VDRC) and the Developmental Studies Hybridoma Bank (DSHB) for reagents, fly stocks and antibodies. Hsi-Yu (Sylvie) Chen for help with midgut dissections. This work was funded by the National Institute of General Medical Science (NIGMS) under award number R35GM118330. The content is solely the responsibility of the authors and does not necessarily represent the official views of the NIH.

## Author contributions

Conceptualization, AB; Methodology, JLL, AA; Formal Analysis, JLL; Investigation, JLL, MT, AA, AS; Resources, AB; Writing, AB; Supervision, AB; Funding Acquisition, AB.

## Materials and Methods

### *Drosophila* husbandry

All crosses were performed on standard cornmeal-molasses medium (60g/L cornmeal, 60ml/L molasses, 23.5g/L bakers yeast, 6.5g/L agar, 4ml/L acid mix and 0.13% Tegosept). Crosses not involving conditional expression of transgenes were incubated at room temperature. Crosses involving conditional expression of transgenes including RNAi were incubated at l 8°C until adult offspring eclosed. Adults were kept at l 8°C for 5 days before they were incubated at 29°C for another 5-6 days prior to dissection. Flies were provided fresh, yeasted food every day. Only female midguts were dissected and analysed. SMURF assays were performed as described ^27^ except flies were incubated on blue food for 18 hours.

### Fly stocks and genetics

*Canton* S (BL64349) and *luciferase* RNAi (BL31603) were used as control stocks. The following *dronc* stocks were used: *dronc*^*I24*^ and *dronc*^*I29* 11^; *UAS-dronc*^*wt*^ and *UAS-dronc*^*C318A* 49^; *UAS-dronc* RNAi: P{GD12376}v23033 and P{KK104278}v100424 from VDRC. Other stocks used were: *UAS-p35* (BL5072); *UAS-drICE* RNAi (BL32403); *UAS-dark* RNAi (BL33924), *esg-Gal4* ^50^; *Su(H)GBE-Gal4* = *Su(H)-Gal4* ^51^; *NPJ-Gal4* ^52^; *tub-Gal80*^*ts* 53^. The stock *Su(H)GBE-Gal4 UAS-GFP/CyO; tub-Gal80*^*ts*^/*TM2* is a kind gift of Hsi-Yu Chen (Ip lab).

### Dissection and immunolabeling of adult guts

Intact female midguts were dissected using standard protocols ^54^. Primary antibodies were: anti Dronc (1:200; ^55^; a kind gift of Pascal Meier); PH3 (1:2,000; Millipore), Prospero (Pros, 1:20; DSHB; Prospero (MR1A) was deposited to the DSHB by C.Q. Doe); Pdm-1 (1:1,000; ^56^; a kind gift of Yu Cai). DAPI was used to counterstain nuclei. Phalloidin labeling was used to assess the physical properties of the guts. Secondary antibodies were donkey Fab fragments from Jackson ImmunoResearch. If not noted otherwise, region R4ab ^32,33^ in the posterior midgut was imaged and analysed. Images were obtained with a Zeiss LSM 700 confocal microscope, analysed with Zen 2012 imaging software (Carl Zeiss) and processed with Adobe Photoshop CS6.

### Counts of DAPI-, Pdm-1-, Pros- and PH3-positive cells

DAPI-, Pdm-1- and Pros-positive cells were counted manually by detecting signal-positive cells as spots in region R4ab and normalized to areas of 2,500 μm^2^. PH3 counts were performed across the entire intestine. At least three independent experiments for every genotype were performed. Analysis and graph generation was done using GraphPad Prism 8.30. The statistical method used is indicated in the legends to the respective panels.

---

**Supplementary Figure S1.**
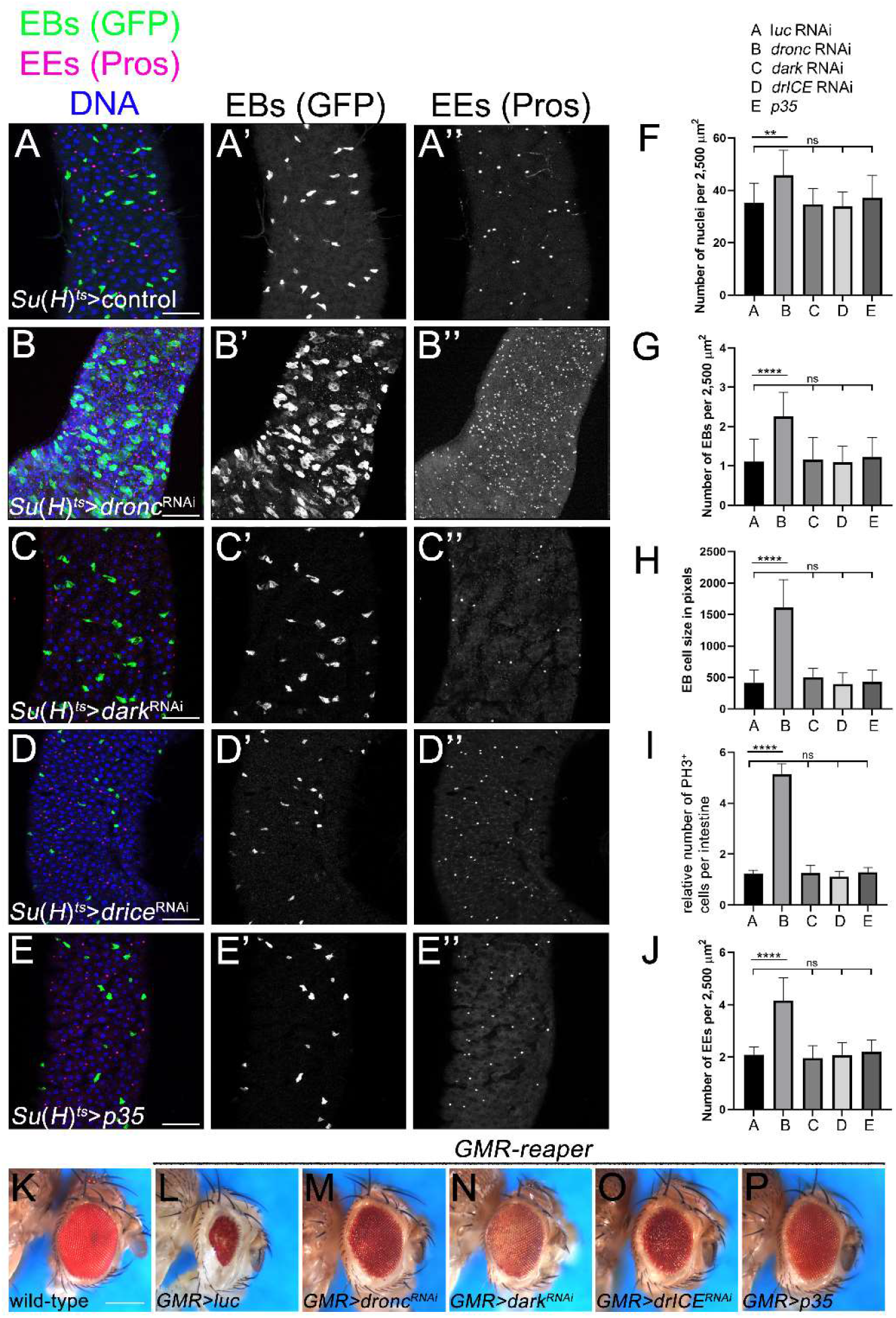
*dark*^RNAi^, *drICE*^RNAi^ and *p35* expression in EBs do not phenocopy the *dronc*^RNAi^ phenotypes in the adult midgut, related to Figure 3. Crosses were incubated at 18°C until adults eclosed. 5 days old females of the indicated genotype were shifted to 29°C and incubated for 5-6 days until dissection and labelling of the midguts. For comparison, the results of *dronc* RNAi are shown in (B) and quantified in (F-J). The *dronc* RNAi transgene used in (B) was P{GD12376}v23033. *Luciferase* RNAi was used as a control in (A). **(A-E)** Shown are R4ab regions of midguts of the indicated genotypes labelled for enteroblasts (EBs) (GFP; green in A-E, grey in A’-E’)), enteroendocrine (EEs) cells (Pros; red in A-E, grey in A’’ -E’’) and nuclei (DAPI; blue in A-E). Scale bar: 50 μm. Complete genotypes: *Su(H)-Gal4 UAS-GFP tub-Gal80*^*ts*^; *UAS*-X (X = *UAS-luciferase* RNAi (A), *UAS-dronc* RNAi (B), *UAS-dark* RNAi (C), *UAS-drICE* RNAi (D), *UAS-p35* (E). **(F-1)** Quantification of the numbers of nuclei (F), EBs (G), EB cell size (H), PH3 counts (I) and number of EEs (J) of the midguts in (A-E). To obtain nuclei counts, at least five random, but representative fields of 2,500 μm^2^ each per posterior midgut (region R4ab) were counted. In the case of EBs and EEs, all GFP-positive cells (corresponding to EBs) and Pros-positive cells (EEs) were counted in region R4ab and normalized to 2,500 μm^2^ fields. For EB cell size calculation, the shape of EBs was outlined using the quick selection tool in Photoshop and the number of pixels in the selected area was recorded. PH3 counts were performed across the entire intestine. Data were analysed by one-way ANOVA with Holm-Sidak test for multiple comparisons. Error bars are SEM. ****p < 0.0001; **p< 0.01. ns-not significant. Number of midguts in (F,G,J) analysed: n=12 (control), 9 (*dronc* RNAi), 10 (*dark* RNAi), 11 (*drICE* RNAi), 10 (*p35)*. For EB cell size calculation (H): n=19 (control), 13 (*dronc* RNAi), 34 (*dark* RNAi), 25 (*drICE* RNAi), 18 (*p35)*. Number of intestines for PH3 counts (I): n=9 (control), 7 (*dronc* RNAi), 8 (*dark* RNAi), 9 (*drICE* RNAi), 7 (*p35)*. Exact genotypes as in (A-E). **(K-P)** Validation of the RNAi and *p35* transgenic lines used in (A-E). The RNAi and *p35* lines used in (A-E) were tested in the apoptosis model *GMR-reaper* which causes an eye ablation phenotype (L) compared to wild-type (Canton S) (K). The *dronc, dark* and *drICE* RNAi lines as well as the *p35* line driven by *GMR-Gal4* suppressed the small eye phenotype of *GMR-reaper* (M-P) suggesting that these transgenes are functional. Exact genotypes: (K) wild-type (Canton S); (L-P) *GMR-reaper GMR-Gal4/UAS-X* (X = *UAS-luciferase* RNAi (L), *UAS-dronc* RNAi (M), *UAS-dark* RNAi (N), *UAS-drICE* RNAi (O), *UAS-p35* (P).

